# Recombinant Human TSH Fails to Induce the Proliferation and Migration of Papillary Thyroid Carcinoma Cell Lines

**DOI:** 10.1101/2024.04.02.587718

**Authors:** Georgios Kalampounias, Athina Varemmenou, Christos Aronis, Irene Mamali, Athanasios-Nasir Shaukat, Dionysios V Chartoumpekis, Panagiotis Katsoris, Marina Michalaki

**Author notes:** **Correspondence:** Panagiotis Katsoris (P.K.). These authors share the last authorship (Panagiotis Katsoris, Marina Michalaki).

## Abstract

Thyrotropin (TSH) suppression is required in the management of patients with papillary thyroid carcinoma (PTC) to improve their outcomes, inevitably causing iatrogenic thyrotoxicosis. Nevertheless, the evidence supporting this practice remains limited and weak, and *in vitro* studies examining the mitogenic effects of TSH in cancerous cells used supraphysiological doses of bovine TSH, which produced conflicting results. Our study explores for the first time the impact of human recombinant thyrotropin (rh-TSH) on human PTC cell lines (K1 and TPC-1) that were transformed to overexpress the thyrotropin receptor (TSHR). The cells were treated with escalating doses of rh-TSH under various conditions, such as the presence or absence of insulin. The expression levels of *TSHR* and *thyroglobulin* (*Tg*) were determined, and subsequently, the proliferation and migration of both transformed and non-transformed cells were assessed. Under the conditions employed, rh-TSH was not adequate to induce either the proliferation or the migration rate of the cells, while *Tg* expression was increased. Our experiments indicate that clinically relevant concentrations of rh-TSH cannot induce proliferation and migration in PTC cell lines, even after overexpression of TSHR. Further research is warranted to dissect the underlying molecular mechanisms, and these results could translate into better management of PTC patients.

## 1 Introduction

Thyroid cancer is the most common endocrine malignancy, accounting for 88% of all endocrine carcinomas and 3% of all human cancers [1–3]. Papillary thyroid carcinoma (PTC) is the most common type of thyroid cancer, accounting for 84% of all cases [4]. In the management of PTC patients, a degree of serum thyroid stimulating hormone (TSH) suppression is required depending on disease severity and response to therapy [1,5]. Thus, a substantial number of PTC patients are subjected to iatrogenic subclinical hyperthyroidism, which is known to have adverse effects on cardiac structure and function as well as bone health [6–9]. The beneficial effects of suppressed TSH on the outcomes of PTC patients are not well-established [10]. In one prospectively followed multi-institutional registry study from the National Thyroid Cancer Treatment Cooperative Study Group in North America on 2,936 differentiated thyroid carcinoma (DTC) patients of all stages, it was found that TSH suppression improves overall survival (OS) and disease-specific survival (DSS) only in high-risk patients [11]. Additionally, maintaining a subnormal to undetectable TSH range in stage II patients has a beneficial effect on OS [11]. On the contrary, in one randomized controlled trial that included 441 patients with DTC of all stages, disease-free survival (DFS) was not more than 10% inferior to those without TSH suppression [12]. In a very recently published systematic review and meta-analysis, the beneficial effects of suppressed serum TSH in intermediate- and high-risk patients are questioned. Nine studies were selected, and in seven of them, progression-free survival (PFS), DFS, and relapse-free survival (RLFS) were similar among patients with suppressed or not suppressed TSH. In four of them, OS and DSS composite outcomes did not differ between the two groups of patients [13]. It is worth mentioning that there is heterogeneity among studies regarding the definition of TSH suppression, ranging from <0.01 to <0.5 μU/mL or lower than the reference range, whereas ‘not suppressed’ means up to 2 μU/mL. The rationale for TSH suppression in the management of PTC patients comes from the notion that TSH, via its receptor, expressed in the follicular epithelial cells of the thyroid gland, regulates thyroid function and growth [14]. Thyroid stimulating hormone receptor (TSHR) couples to G proteins of all four subfamilies, including the stimulatory G protein Gs, which activates adenylyl cyclase to produce cyclic adenosine monophosphate (cAMP); Gi, which inhibits cAMP production; G13, which activates p44/42 mitogen-activated protein kinases (MAPKs - commonly known as ERK1/2); and Gq/G11, which activates phospholipase C to produce inositol-1,4,5-trisphosphate, which is rapidly degraded to inositol monophosphate (IP-1; phosphoinositide signaling). From another perspective, activating mutations of either *TSHR* or *Ga* proteins that underlie functioning thyroid nodules preclude thyroid cancer [1,15]. Similarly, autoantibodies against TSHR in Graves’ disease are not widely recognized as a risk factor for thyroid cancer development [15,16]. Nevertheless, intriguing findings from a mouse model carrying a patient-derived constitutively active *TSHR D633H* mutation reveal a different scenario [17]; large papillary thyroid tumors emerged in mice carrying the mutation at around one year of age, with nearly all homozygous mice affected. Remarkably, the commonly observed driver mutations of *Braf*, *Nras*, and *Kras* were notably absent in these tumors. Thus, it seems that enhanced TSHR signaling due to some rare *TSHR* activating mutations could lead to thyroid cancer development and growth. However, in the most common thyroid cancers due to *Braf*, *Nras*, and *Kras* mutations, the role of TSHR signaling in the development and/or growth of these tumors is not extensively studied. Hence, the role of TSH suppression in all these cases warrants investigation.

Few *in vitro* studies of various thyroid cell culture systems treated with different doses of cattle/bovine TSH (b-TSH) give inconclusive results regarding the proliferation and function of thyroid cells [18–23]. The doses of TSH used in these studies were extremely high compared to circulating concentrations in humans. Recently, recombinant human thyrotropin (rh-TSH) became available on the market. Human TSH has 88% homology with its bovine counterpart (NCBI Homologene). Bovine TSH contains positively charged amino acid residues at the α-subunit, that are known to increase hormone-binding properties compared to human TSH [24]. The biological and immunological activities of rh-TSH were examined in human fetal thyroid cells [25]. Its biopotency was estimated to be 4.3±0.7 IU/mg and the immunopotency to 10.2±0.7 IU/mg whereas the immunopotency of b-TSH is 20-40 IU/mg and highly dependent on purification [26].

Herein, we investigated for the first time the effects of rh-TSH in physiologically relevant doses resembling the TSH concentrations in human plasma on the function and proliferation of PTC human thyroid cell lines. For this purpose, we selected two well established cell lines, K1 and TPC-1, both of which adequately model PTC cells and express the main thyroid markers, thus allowing their classification as PTC cells and indicting their differentiation status [27,28]. To fully assess the role of rh-TSH in PTC cell proliferation, not only did we co-administer it with recombinant human insulin (rh-INS), but we also created cells that controllably overexpress the TSH receptor. TSHR is known to be a key protein in the signal transduction pathways that control TSH-dependent cell proliferation, migration and differentiation. Nonetheless, in most *in vitro* settings, *TSHR* expression is significantly downregulated, and to multiply the incoming mitogenic signals, we used an inducible gene expression system to overexpress the receptor and assess its role.

## 2 Materials and Methods

### 2.1 Cell Lines and Culture Conditions

The K1 and TPC-1 PTC cell lines were used as models for our study (ECACC, Salisbury, UK). K1 were cultured in high-glucose DMEM (Biowest, Nuaille, France), while TPC-1 cells were grown in RPMI 1640 (Biowest, Nuaille, France), respectively, supplemented with 10% fetal bovine serum (FBS) (Biowest, Nuaille, France) and 1% penicillin/streptomycin (Biosera, Nuaille, France) in a 37°C incubator with 5% CO_2_. All cell culture expendables were purchased from Greiner Bio-One (Kremsmünster, Austria).

### 2.2 Thyroglobulin (Tg) and TSHR Expression Determination Using Quantitative Polymerase Chain Reaction (qRT-PCR)

To assess the baseline expression levels of *Thyroglobulin* (*Tg)* and *TSHR*, as well as *Tg* induction as a result of rh-TSH stimulation in K1 and TPC-1 cells, qRT-PCR was used. Cells were treated with designated rh-TSH doses, and 24 h later, total RNA was extracted using the Monarch® Total RNA Miniprep Kit (New England Biolabs, Ipswich, MA, USA). The extracted RNA was eluted in 50 μL of nuclease-free water and treated with Qiagen DNAse I (Germantown, MD, USA), following the manufacturer’s procedure. The RNA quantity was assessed using the NanoDrop Q500 spectrophotometer (Quawell CF, USA). RNA quality was assessed by determination of the ratio for absorbance at 260 nm vs. absorbance at 280 nm (A260 nm/A280 nm) using the NanoDrop Q500 spectrophotometer (which was assessed by calculating the ratios of absorbance A260nm/A280nm and A260nm/A230nm). RNA integrity was roughly verified by electrophoresis on a 1.2% denaturing agarose gel stained with ethidium bromide. 1μg of total RNA was reverse transcribed using the Transcriptor First Strand cDNA Synthesis Kit (Roche Life Sciences, Penzberg, Germany), according to the manufacturer’s instructions. The cDNAs were subjected to quantitative PCR (qRT-PCR), which was performed with the LightCycler® 2.0 Real-Time PCR System (Roche Life Sciences, Penzberg, Germany) using LightCycler® FastStart DNA Master SYBR® Green I (Roche Life Sciences, Penzberg, Germany). The cycling conditions consisted of 15 minutes of denaturation at 95 °C, followed by 45 cycles of 95 °C for 10 sec 60 °C for 30 sec, and 72 °C for 20 sec. The primers used in qRT-PCR were for *Tg* mRNA detection:

*Tg* forward 5΄-CACCAACTCCCAACTTTTCC-3΄

*Tg* reverse 5΄-CAACTGACCTCCTTTGCCA-3΄

The primers used for *TSHR* mRNA expression were:

*TSHR* forward 5’-GGAATGGGGTGTTCGTCTCC-3’

*TSHR* reverse 5’-GCGTTGAATATCCTTGCAGGT-3’

As a reference gene, *ACTB* (*β-actin*) was used, which was detected with the following primers:

β-actin forward 5΄-GCACAGAGCCTCGCCTT-3΄

β-actin reverse 5΄-GTTGTCGACGACGAGCG-3΄

All primers were purchased from IDT (Leuven, Belgium). The DNA polymerase FIREPol® and the mastermix were from Solis BioDyne (Tartu, Estonia). The specificity of the PCR product was verified by melting curve analysis (single-peak). For qRT-PCR analysis, the data were analyzed using the relative quantification method. The efficiency of each PCR reaction was calculated by serial dilutions of pooled cDNA samples [29].

### 2.3 Proliferation Assays of K1 and TPC-1 Cells

#### 2.3.1 Treatment with Recombinant Human TSH and/or Recombinant Human Insulin

Equal numbers of K1 or TPC-1 cells (25, 000) were seeded on 48-well TC-treated plates (Greiner Bio-One, Kremsmünster, Austria) and left for 24 h to settle. Then, the cells were rinsed twice with a PBS solution, and treated with recombinant human TSH (rh-TSH) in the form of Thyrogen® (Sanofi Genzyme, Cambridge, MA, USA) and/or recombinant human insulin (rh-INS) in the form of Humalog® (Eli Lilly and Company, Indianapolis, IN, USA). Increasing concentrations were used (as indicated in the relevant figures) in the presence of 2.5% FBS for 48 or 72 hours. (5-1,000 μIU/mL). The 2.5% FBS concentration was chosen based on preliminary experiments as the minimum (from a range of 1–10%) that allows the cells to survive and replicate as expected. Simultaneously, using a minimal FBS concentration was crucial to avoid stimulation of the cells by significant amounts of hormones contained in the bovine serum (endogenous b-TSH, bovine insulin-like growth factor 1 (b-IGF1), insulin (b-INS), and relative co-factors), since the purpose was to administer controlled doses of TSH and insulin. A negative control was included in each experiment (culture medium supplemented with 2.5% FBS, without additional rh-TSH/rh-INS) as well as a positive control with an elevated FBS concentration of 10%. FBS could not have been omitted due to the thyroid cells need for growth factors and iodine salts [30]. To maintain the thyroidal characteristics of the cells (avoid de-differentiation), allow them to proliferate (as all the aforementioned factors are needed), and suppress apoptosis signals caused by the lack of proper hormonal stimulation, this minimal FBS concentration was necessary and thus maintained during all assays. A detailed description of the FBS concentration selection is presented in Appendix A.

#### 2.3.2 Crystal Violet Assay

Following treatment with the hormones (rh-TSH, rh-INS) for 48-72 h, the final number of viable cells was calculated using the crystal violet assay [31]. The culture media were aspirated, the cells were rinsed twice with a phosphate buffered saline (PBS) (pH = 7.4) solution, and then fixed for 10 minutes at 25 °C in a 3.7% formaldehyde solution in PBS. Then, the fixation was aspirated, the cells were rinsed, and subsequently stained in a methanolic crystal violet solution (Sigma-Aldrich, Burlington, MA, USA) for 20 minutes at 25 °C. Following staining, the cells were rinsed three times with distilled water to remove any residual stain. Then, the water was carefully aspirated, and a 30% acetic acid solution was added to extract the cell-bound crystal violet dye by mild shaking for 20 minutes at 25 °C. Finally, the extracted solution was transferred to a 96-well flat bottom plate (Greiner Bio-One, Kremsmünster, Austria) for spectrophotometry at 590 nm using a microplate reader. The equation used to convert the optical density (OD) to a number of cells (N) was the following: N = 0.5 x 10^5^ x OD. This equation was based on preliminary experiments with an increasing number of cells stained with the above-mentioned method.

### 2.4 Wound Healing Assays of K1 and TPC-1 Cells

The wound healing assay was used as a rapid way to assess the hormones’ effects on the migratory potential of both K1 and TPC-1 cells and has also been employed by other researchers [32,33]. Cells were seeded in a 6-well plate (Greiner Bio-One, Kremsmünster, Austria) until the formation of a complete monolayer. A scratch was then formed in the cells’ monolayer, and after washing away the scraped cells with a PBS solution, the cells were incubated with rh-TSH concentrations of 5, 10, 20, 50, and 100 μIU/mL and 2.5% FBS for 72 hours and 0.5 U/mL rh-INS. Consequently, the healing rate was estimated by capturing images of the wounds at key time points (24, 48, and 72 h) using an inverted microscope.

### 2.5 Creation of the K1-TSHR and TPC-1-TSHR Cell Lines Using the Lenti-X Tet-on Advanced System

To multiply the TSH signal reception, cell clones were created to overexpress the TSHR receptor in a controllable manner. Instead of using mutant types of the *TSHR* gene (many of which have been reported), the wild type allele was selected since most *TSHR* mutations are associated with follicular thyroid carcinoma (FTC) much more often compared to PTC, where mutations are relatively rare [34]. Additionally, wild-type TSHR (*wt-TSHR*) has been reported to possess a high level of constitutive activity in the wild-type form [35]; thus, overexpressing would significantly multiply signal reception and transduction.

#### 2.5.1 Transduction of K1 and TPC-1 Cells Using Lentiviral Vectors

Human *wt-TSHR* cDNA was subcloned from a pcDNA 3.1 hygro (-) plasmid (as a kind gift from Dr. Holger Jaeschke from University Hospital Essen, Germany) to a pLVX-Tight-Puro plasmid [36]. This plasmid was used to transfect HEK293T cells (ECACC, Salisbury, UK) to produce lentivirus. Plasmids containing the Lenti-X Tet-on (pLVX-Tight-Puro) advanced system and the embedded cDNA *TSHR* sequence were constructed using the restriction enzymes BamHI and EcoRI. Subsequently, the transformed plasmids were analyzed using agarose electrophoresis, and the constructed plasmid was isolated and transferred to *Escherichia coli* DH5a bacteria. The *E. coli* bacteria were cultured in an LB growth medium containing 100 μg/mL ampicillin; thus, only the transformed clones survived. Plasmids were then isolated using the Monarch® Plasmid Miniprep Kit (New England Biolabs, Ipswich, MA, USA) and used to transfect HEK293T cells. The culture supernatant containing the viruses produced in the HEK293T cells was used to transfect K1 and TPC-1 cells. Following a 24-hour incubation with the lentiviruses, the medium was replaced with puromycin- and G418-containing medium at concentrations of 150 μg/mL. After 48 hours, only the transformed cells had survived and were still attached.

#### 2.5.2 Transformation Validation Using Polymerase Chain Reaction

DNA was extracted from lentivirus-transduced cells (K1 and TPC-1) using the Monarch® Genomic DNA Purification Kit (New England Biolabs, Ipswich, MA, USA), and its quality was determined using the NanoDrop Q500 spectrophotometer (Quawell CF, USA). PCR was performed to detect the TSHR inserts using the primers:

TSHR forward 5’-ATGGGACAAAGCTGGATGCT-3’ TSHR reverse 5’-AGCAAGCTTGGTCCACTGTA-3’

The housekeeping gene *ACTB* (β-actin) was used as a reference mRNA. All primers were purchased from IDT (Leuven, Belgium). The cycling conditions consisted of 3 minutes denaturation at 95 °C, followed by 30 cycles of 95 °C for 3 sec, 60 °C for 30 sec and 72 °C for 30 sec. The DNA polymerase FIREPol® and the mastermix were from Solis BioDyne (Tartu, Estonia).

### 2.6 TSHR mRNA Overexpression Validation in K1-TSHR and TPC-1-TSHR cells

#### 2.6.1 Stimulation with Doxycycline and Quantitative Polymerase Chain Reaction (qRT-PCR)

To assess the induction of *TSHR* mRNA expression in both transformed cell lines (they were named K1-TSHR and TPC-1-TSHR, respectively) a dose of 3 μg/mL doxycycline (DOX) was administered, and the procedure described in paragraph 2.2 was followed. The DOX dose was selected following optimization of the system as described in previous publications [37]. Non-transformed cells (K1, TPC-1) were also monitored as a control group.

#### 2.6.2 Receptor Synthesis Validation with Western Immunoblotting

To assess the TSHR protein levels in both transformed (K1-TSHR and TPC-1-TSHR) and non-transformed (K1 and TPC-1) cells, crude lysates were prepared using Laemmli Sample Buffer and were subsequently loaded onto 12% polyacrylamide gels for SDS-PAGE analysis. The proteins were then transferred onto the 0.20 μm PVDF Porablot® membrane (MN, Nordrhein-Westfalen, Germany), and Western blot (WB) analysis was performed using an Anti-TSH Receptor/ TSHR-R rabbit polyclonal IgG antibody (ab58917) (Abcam, Cambridge, UK). The detection was performed using a secondary Anti-rabbit IgG HRP-linked goat antibody (CST #7074) and the chemiluminescence system SignalFire™ ECL (CST #6883) both from Cell Signaling Technology (Danvers, MA, USA). As a loading control, the cytoskeletal protein α-tubulin was used, which was detected using an Anti-Tubulin mouse monoclonal (6-11B-1) IgG2b antibody (#T6793) (Sigma-Aldrich, Darmstadt, Germany), and a secondary Anti-mouse IgG HRP-linked horse antibody (CST #7076) from Cell Signaling Technology (Danvers, MA, USA). As a housekeeping gene, α-tubulin had been found to remain stably expressed during all treatments, and its accumulation was relatively similar among the different cell clones studied. The normalization was performed using a plugin for ImageJ created by Suarez-Arnedo (2020) [38].

### 2.7 Proliferation and Migration Assays of K1-TSHR and TPC-1-TSHR Cells

To determine alterations in the proliferation rate descending from TSHR overexpression, the cells were stimulated with doxycycline for 24 h to synthesize an excessive amount of the receptor, as had been demonstrated using qRT-PCR and WB. During the initial 24-hour settling period of the proliferation assays, doxycycline was present (at a concentration of 3 μg/mL) to achieve TSHR overexpression. Following the incubation procedure with the hormones (as described in paragraph 2.3), the cell number was estimated using the crystal violet assay. To assess possible interactions between doxycycline and the hormones, two sets of control experiments were performed: a) non-transformed cells (K1, TPC-1) were exposed to doxycycline in the same way as transformed cells (K1-TSHR, TPC-1-TSHR); b) and transformed cells were also assessed in the absence of a doxycycline pre-treatment, theoretically functioning as naïve K1/TPC-1 cells.

### 2.8 Statistical Analysis

To compare the number of cells among the different concentrations of rh-TSH or rh-INS, a one-way ANOVA test was used, followed by Sidak’s multiple comparisons test. The results are displayed as means ± the standard error of the mean (SEM). Prism 8 (Graphpad, La Jolla, CF, USA) for Windows was used for the statistics test and the generation of graphs. Statistical significance was set at p-value<0.05.

## 3 Results

### 3.1 Effects of escalating rh-TSH concentrations on Thyroglobulin Expression, Proliferation and Migration Rates of K1 and TPC-1 Cells

Both cancerous cell lines, under the culture conditions applied expressed low levels of *Tg* mRNA which were quantified using qRT-PCR. The transcription of the *Tg* gene was used as marker of the thyroidal phenotype maintenance, and any differences of the *Tg* expression were theorized as results of rh-TSH and insulin administration. Since both cell lines synthesized low levels of Tg, the detection of the thyroglobulin protein using western blots or ELISA kits was not feasible, and we thus settled on qRT-PCR results. The low expression was verified by the high C_t_ numbers observed in all qRT-PCR experiments. K1 cells were found to synthesize slightly lower *Tg* compared to TPC-1 cells; nonetheless, both cell lines’ expression was low compared to bibliographically mentioned information about tumors from PTC patients, where *Tg* levels are comparable to those of normal thyroid tissue [39]. To study the cells’ response to rh-TSH stimulation, we incubated them with designated doses of the hormone to assess their activity regarding *Tg* synthesis. Stimulation with rh-TSH (10 μIU/mL, 100 μIU/mL) led to an increase in *Tg* mRNA expression of both K1 and TPC-1 cells (**Figure 1a**) (p-value<0.01), indicating that rh-TSH indeed activates the thyroid-specific signaling pathways that lead to *Tg* expression.

**Figure 1.**
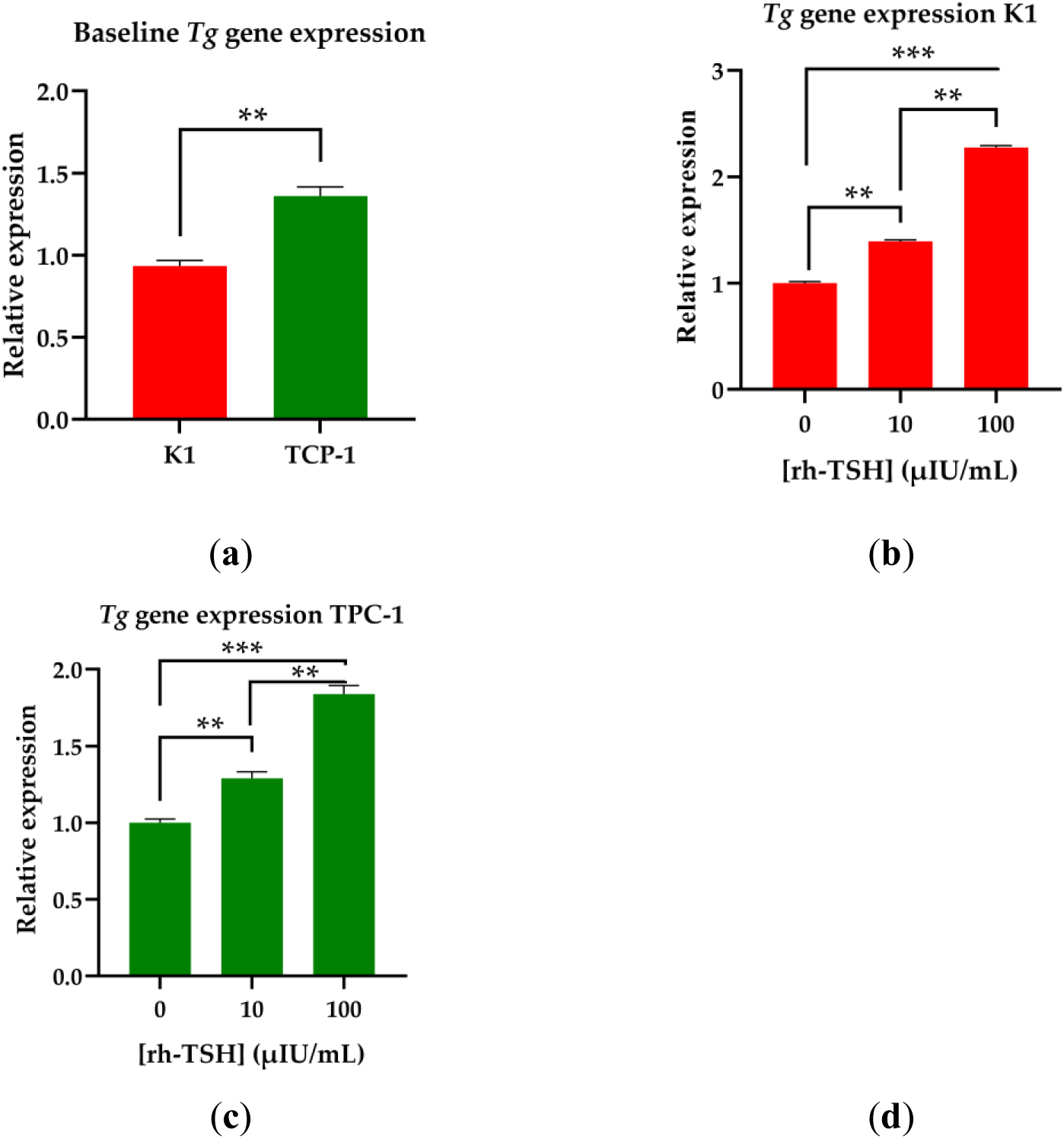
*Thyroglobulin* (*Tg*) Gene Expression Determination. **(a)** Using qRT-PCR, the baseline expression levels of the *Tg* mRNA of K1 (red bar) and TPC-1 cells (green bar) were assessed. **(b)** Following 48 h of treatment with the designated doses of rh-TSH, qRT-PCR was used to assess the effects of the hormone on *Tg* mRNA expression in K1 cells, while the same assay was performed in TPC-1 cells as well **(c)**. All the samples were analyzed using Livak’s model, and beta-actin was used as a reference gene to normalize the data. All experiments were conducted in triplicate and each bar represents the average of the three experimental values. The error bars refer to the standard error of the mean (SEM). * Corresponds to a p-value < 0.05; ** corresponds to a p-value < 0.01; *** corresponds to a p-value <0.001.

Regarding proliferation, K1 cells were incubated for 48 hours with increasing concentrations of rh-TSH of 0, 1, 5, 10, 20, 50, and 100 μIU/mL in high-glucose DMEM (**Figure 2a**). A concentration of 2.5% FBS was used in all the proliferation experiments, as it was determined to be the minimum dose needed for sustaining cell survival and proliferation, therefore avoiding possible masking effects of the rh-TSH action by serum-contained bovine TSH (b-TSH) or other hormones (**Figures A1a and A1b**). Iodine intake has been found to be a limiting factor in thyroid cell proliferation (K1 and TPC-1) in *in vitro* settings [30]; therefore, it could not be absent. According to our data, no discernible differences in cell number were documented between the various rh-TSH concentrations; therefore, a range of higher concentrations between 100 and 1,000 μIU/mL was tested (**Figure 2b**). The action of rhTSH was examined under various conditions, such as the presence or absence of insulin as shown in all graphs, a prolonged exposure to the hormones (72 h) (**Figure 1c**), and the replenishment of the cell medium every 24 hours (to supply the cells with an abundance of rh-TSH) (**Figure 2d**). Even under those conditions, rh-TSH was incapable of significantly inducing proliferation compared to the control group, which consisted of cells cultured with 10% FBS. This group had elevated proliferation rates compared to the rest of the samples and this activity was a direct consequence of increased serum levels, which contains adequate quantities of pro-proliferative molecules to sustain continuous mitoses, mainly IGFs and iodine salts. The same screening conditions were used to monitor the cancerous cell line TPC-1 (**Figures 2e-2h**), and the results showed that rh-TSH failed to induce cell proliferation in any tested concentration (0–1,000 μIU/mL) regardless of the presence of insulin. All insulin-treated samples showed significantly elevated cell numbers compared to the insulin-untreated group, whereas the rh-TSH concentration did not show an additive effect on cell number (**Figure 2**).

**Figure 2.**
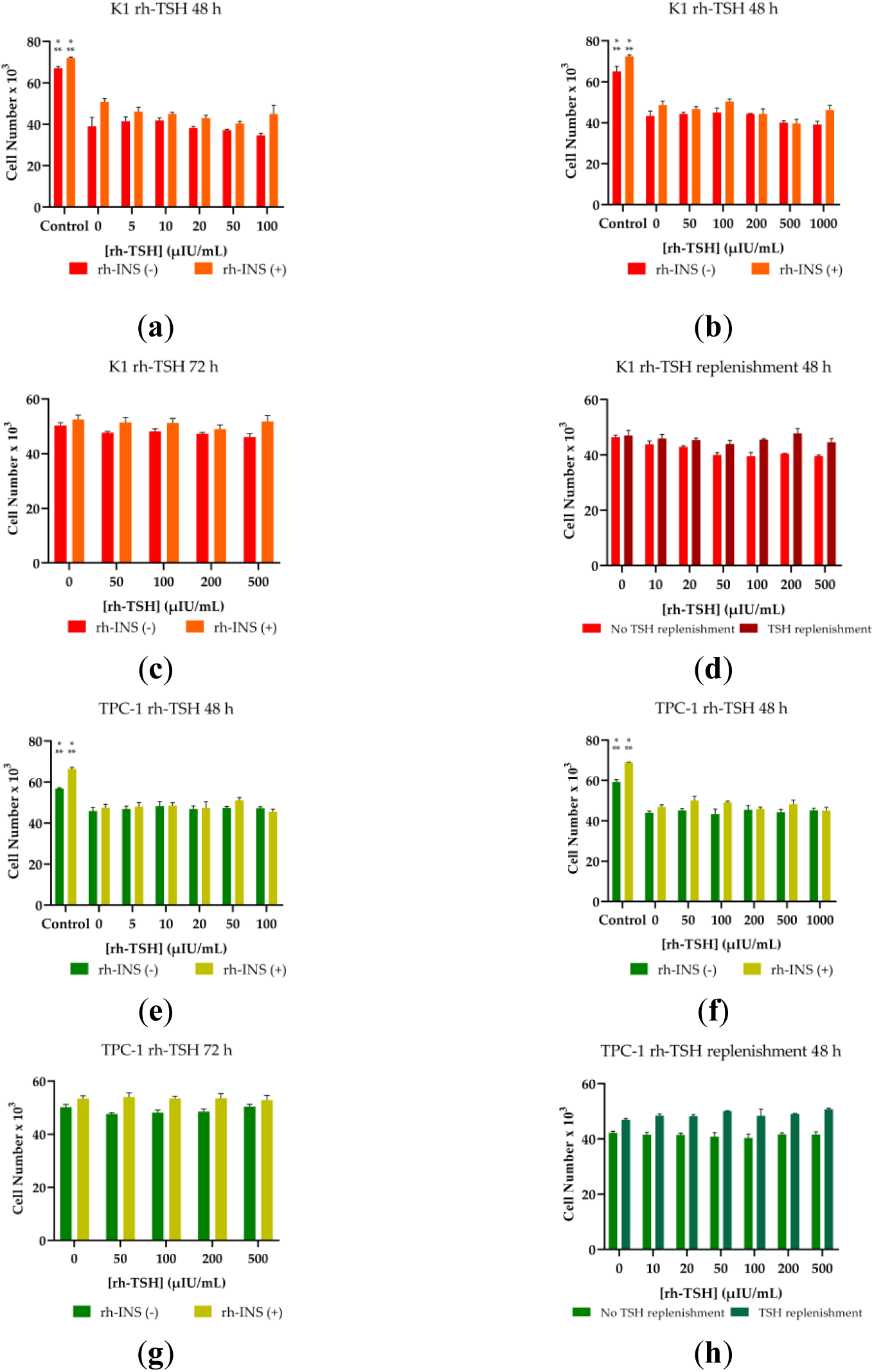
Proliferation Assay of K1 and TPC-1 Cells Treated With rh-TSH and/or rh-INS. Cells were seeded into microplates, and after 24 hours of settling and attachment, the old medium was aspirated and replaced with a new one containing 2.5% FBS and the designated doses of the hormones (rh-TSH, rh-INS). The red bars represent rh-INS-untreated K1 cells, while the orange bars represent rh-INS-treated K1 cells. (**a**) A range of rh-TSH concentrations from 5 μIU/mL to 100 μIU/mL was initially tested in K1 cells, (**b**) as well as a broader spectrum from 50 μIU/mL to 1000 μIU/mL. (**c**) Assays were performed over a 72-hour timeframe. (**d**) To test whether TSHR was rapidly absorbed by the cells/decomposed or inactivated, we replenished the hormone-containing medium after 24 h and let the cells grow. (**e-h**) The aforementioned conditions were replicated on the TPC-1 cells line; The dark green bars represent rh-INS-untreated TPC-1 cells, while the light green bars represent rh-INS-treated TPC-1 cells. Wherever insulin is annotated (rh-INS), a concentration of 0.5 U/mL is used. As a positive control for elevated cell proliferation rates, a 10% FBS-containing medium was used which has endogenous b-TSH,b-IGF-II, insulin and other cofactors. All experiments were conducted in triplicate, and the error bars represent the standard error of the mean (SEM). The data analysis was performed using multiple t-tests, and no significant differences between the different rh-TSH concentrations were detected. Insulin-treated cells exhibited an overall increased cell number, which was statistically significant as shown by a paired t-test (p-value<0.001). * Corresponds to a p-value < 0.05; ** corresponds to a p-value < 0.01; *** corresponds to a p-value <0.001.

Regarding the effects of rh-TSH/insulin on cancer cell migration, the wound healing assay was used. The results showed that rh-TSH was incapable of significantly inducing migratory effects on any of the two PTC cell lines, whether insulin was present or not (**Figures 3a and 3b).** The assays were performed in a 72-hour timeframe, and FBS was present in all samples, since in its absence, the cells do not migrate or multiply to cover wounds. However, the concentration of the serum was maintained at a minimum (2.5%) to reduce the masking effects on rh-TSH. Lower FBS concentrations (1%) have been used by other groups in cell cultures of K1 and TPC-1 cells [33]; however, in these experiments, the assay lasted 48 h, and no report on the expression of thyroid markers exists. A complete matrix of the images from all concentrations monitored and all time points can be found in the Appendix B section (**Figures A4a and A4b**).

**Figure 3.**
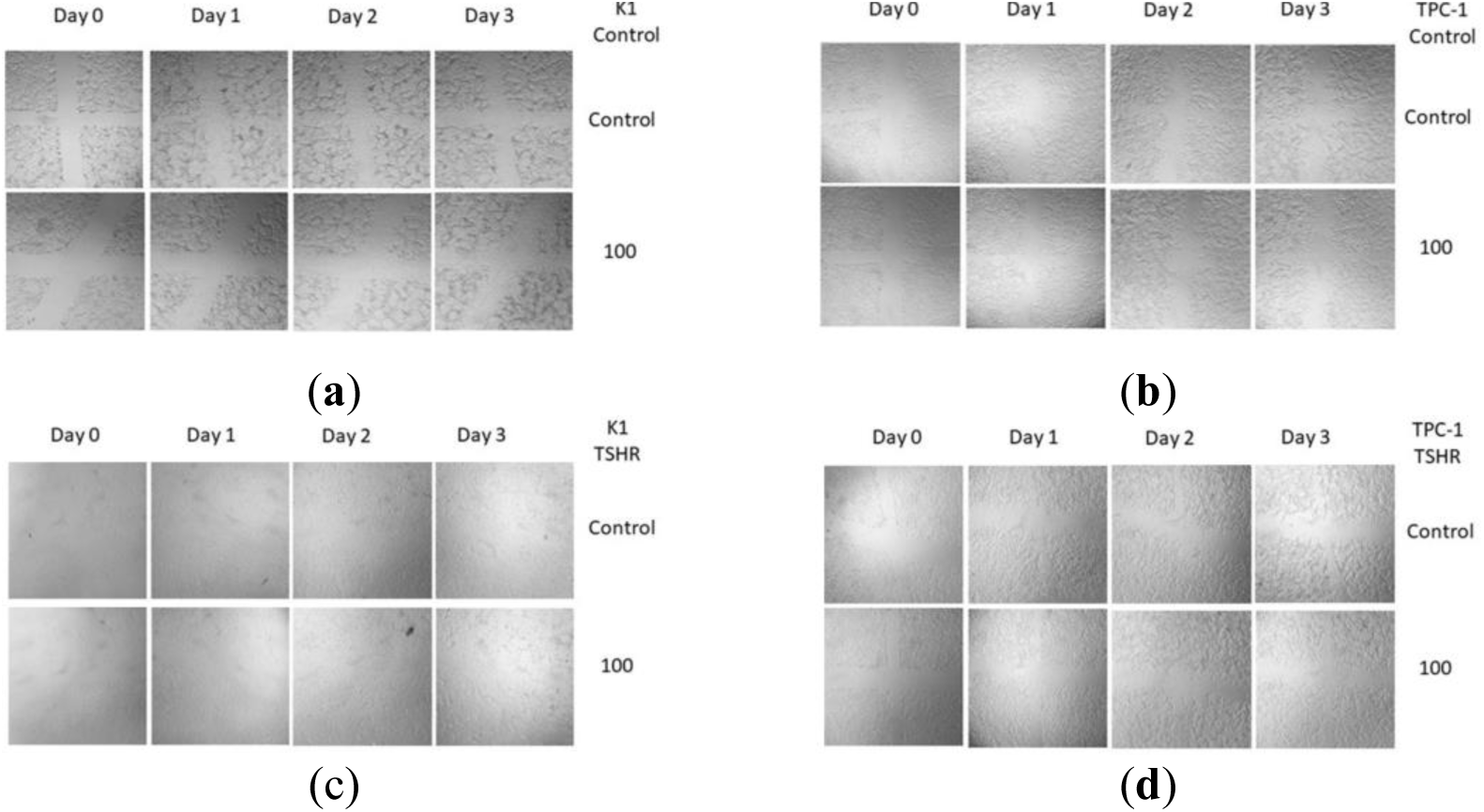
Wound Healing Assay of K1, TPC-1, K1-TSHR, and TPC-1-TSHR Cells Treated With rh-TSH. Cells were seeded into six-well microplates, and after the formation of a confluent monolayer, scratches of equal dimensions were made using a pipette tip. Subsequently, the old medium and the debris were aspirated and replaced with a new one containing 2.5% FBS and the designated doses of the hormones. Photographs were taken every 24 hours using a camera mounted on a microscope (magnification X100). (**a**) Changes were not observed in either K1; or (**b**) TPC-1 cells following treatment with the high dose of 100 μIU/mL, and the same phenomenon was documented on (**c**) K1-TSHR; and (**d**) TPC-1-TSHR cells.

### 3.2 Creation of TSHR-overexpressing PTC cells and TSHR Synthesis Determination

At baseline conditions, qRT-PCR showed that both K1 and TPC-1 cells express low levels of the *TSHR* mRNA. (**Figure 1d**). This low expression has also been verified by other groups [23,27,28,40]; while some others even cite *TSHR* expression as undetectable [41–43]. Low *TSHR* expression allows only for limited stimulation by circulating TSH molecules and this was theorized to be a great limitation of the hormone’s role study. Therefore, to eliminate the problem of low *TSHR* levels, we generated clones of K1 and TPC-1 cells with controllable *wt-TSHR* expression, using the Lenti-X Tet-on Advanced system. The wild-type receptor was selected due to its prevalence in PTC patients and its increased constitutive activity compared to many mutations [34,35]. Plasmids were constructed containing the wild-type *TSHR* gene, and the successful construction was verified with DNA agarose gel electrophoresis (**Figure A2a**). The successful transfection and subsequent transformation were verified using PCR to detect copies of the *TSHR* cDNA insert. Both cell lines had been successfully transformed, and thus the insert was successfully embedded inside the genome (**Figure A2b**). The transformed cells were named K1-TSHR and TPC-1-TSHR, respectively. Following the successful embedment, the expression of TSHR should be inducible through the addition of doxycycline. In order to confirm that, 3 μg/mL of doxycycline (DOX) was introduced for 24 h, and the mRNA levels were assessed by qRT-PCR. The K1-TSHR cells reached an almost 25-fold overexpression, and the TPC-1-TSHR cell line increased its mRNA *TSHR* levels 60 times more than the naïve one (**Figure 4a**). To further confirm that the observed changes at the mRNA level of *TSHR* were indeed translated into differential protein expression, TSHR protein accumulation (after stimulation of both cell lines with DOX) was analyzed by western immunoblotting. The cancerous K1-TSHR overexpressed the receptor by four times and the TPC-1-TSHR cancerous cells eight times, respectively (**Figures 4b-c**).

**Figure 4.**
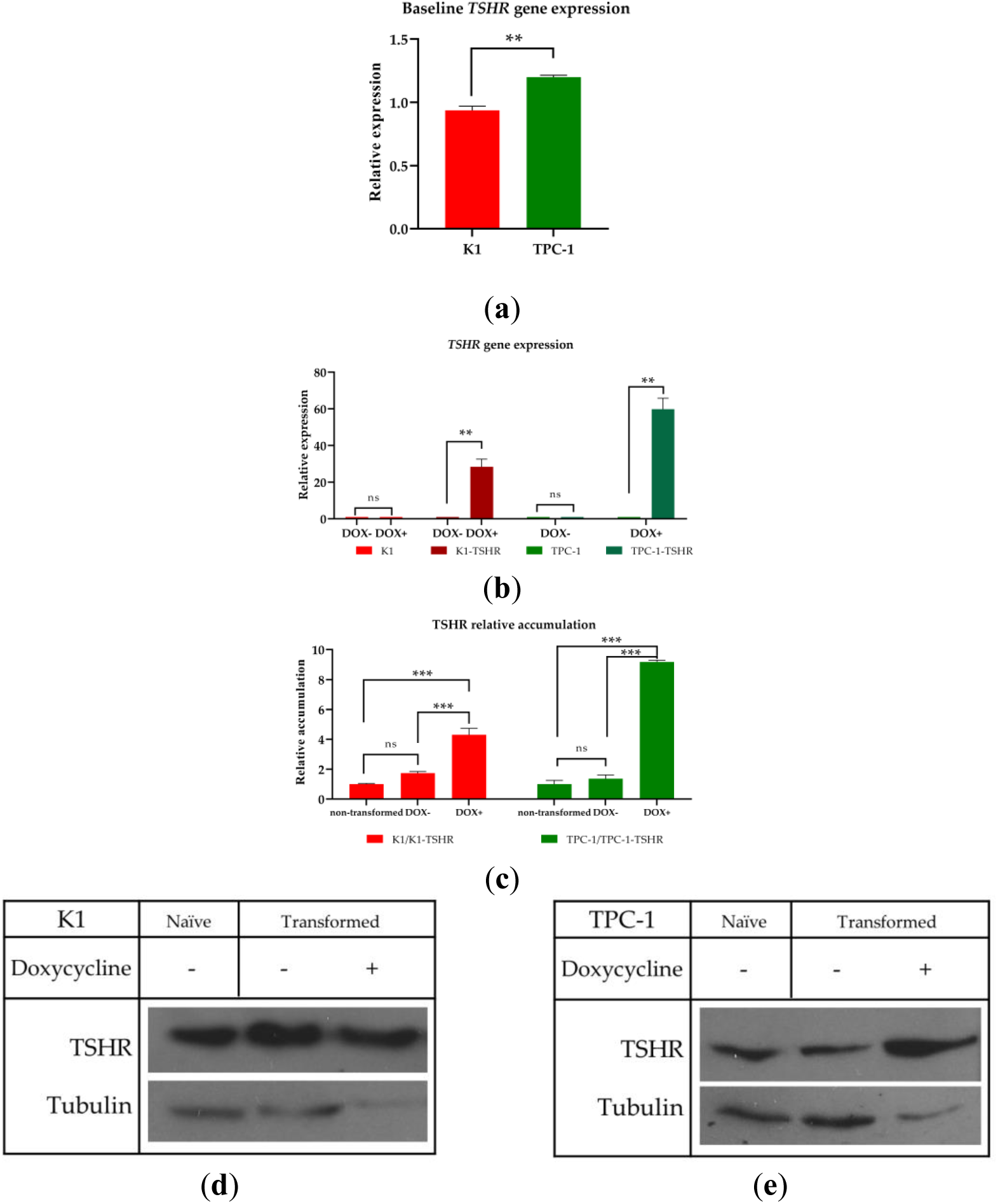
TSHR mRNA and Protein Expression Levels Following Transformation with the Lenti-X Tet-on Advanced System. (**a**) Using qRT-PCR, the baseline expression levels of the *TSHR* mRNA of K1 (red bar) and TPC-1 (green bar) cells were assessed. (**b**) After the selection of the transformed clones using the appropriate antibiotics, incubation with a dose of 3 μg/mL DOX was introduced to the transformed clones to activate the Tet-On Advanced system. Following 24 h of incubation, total RNA was isolated, cDNA was constructed, and qRT-PCR was performed. Relative expression was assessed using Livak’s model. The K1 cell line is annotated with red bars, while the K1-TSHR cells are represented with brown bars. The TPC-1 cell line appears as green bar, while the TPC-1-TSHR cells appear as light blue bars. Each bar represents the average of three experimental values, and the error bars refer to the standard error (SEM). * Corresponds to a p-value < 0.05; ** corresponds to a p-value < 0.01; *** corresponds to a p-value <0.001. Beta-actin was used as a reference gene to normalize the data. (**c**) Following 24 h of incubation with 3 μg/mL DOX, the cells were lyzed, and the crude protein extracts were analyzed using SDS-PAGE on 12% polyacrylamide gels. Western Blot (WB) analysis was performed to detect the TSHR protein band. Using ImageJ, the WB results were quantified (n=3). Representative western blot images are shown of: (**d**) K1; (**e**) and TPC-1. The cytoskeletal protein α-tubulin was used to normalize the results. Due to the increased TSHR synthesis, a lower amount of K1-TSHR/TPC-1-TSHR total protein extract was loaded in the gels to avoid overexposure. The graphs were prepared after normalization with α-tubulin and consideration of the initial difference in protein quantity.

### 3.3 Effects of rh-TSH Concentrations, on Proliferation and Migration of K1-TSHR and TPC-1-TSHR Cells

Proliferation and migration experiments were carried out on transformed K1 and TPC-1 cells after pre-treatment with 3 μg/mL DOX to overexpress the receptor. Comparing the DOX-treated cells to the untreated group, which served as the control group, overall slightly lower cell numbers in the DOX-treated cells were observed (**Figures 4b, 4e**), which were justified as a mild cytostatic consequence of DOX during the overexpression induction period [44]. The same cytostatic effect was observed on the non-transformed cells (**Figures 4c and 4f**). Although the dose used during transduction experiments is far lower than cytotoxic levels, the presence of DOX was expected to mildly affect the proliferation rate, which could be evident due to low FBS concentrations. To guarantee that the effects of rh-TSH were not being masked by DOX, K1-TSHR and TPC-1-TSHR cells were also pre-treated with 3 μg/mL DOX, and their proliferation was assessed. As shown in the figure, similar results were obtained (**Figures 4a and 4d**). The positive control (10% FBS) of the study, regardless of the slightly lower proliferation rate-resulting from DOX-, exhibited successful proliferation which was far greater than that of FBS-starved cells. Therefore, the presence of DOX was not theorized as responsible for rh-TSH ineffectiveness or masking in promoting mitosis since in the case of elevated FBS it did not act as the limiting factor. Insulin treatment (rh-INS) caused all samples to have increased proliferation rates; however, this was observed to be independent from the rh-TSH doses. Even when TSHR was being overexpressed, rh-TSH did not indicate a dose-dependent mode of action when administered as the sole pro-proliferation factor. FBS was found to be the most important parameter, and its effects were further amplified following treatment with insulin.

Regarding migration potential and wound healing capacity, the scratch tests revealed that stimulation with rh-TSH (with or without rh-INS) did not cause any changes (**Figures 3c and 3d**) even when TSHR was overexpressed. A complete matrix of the images from all concentrations monitored and all time points can be found in the Appendix C section (**Figures A4a and A4b**).

## 4 Discussion

Herein, we examine for the first time the effects of clinically relevant concentrations of recombinant human TSH (Thyrogen®) on the proliferation and migration of papillary cancer thyroid cell lines, in contrast with previous studies that mostly employed bovine TSH. Surprisingly, we found that the administration of escalating doses of rh-TSH, like those found in plasma, did not increase cell number or migration rate in K1 and TPC-1 papillary thyroid carcinoma cell lines, which is a rather new notion since only a few groups have come up with similar conclusions. Since TSH signaling significantly depends on the level of thyroid cell (de)differentiation, we assessed the expression of thyroid markers following administration of rh-TSH, and we found that it tended to increase the levels of *Tg* mRNA, a direct proof of successful stimulation. Due to the low expression of TSHR in these cell lines (**Figure 4a**), we successfully transformed them to overexpress TSHR (**Figure 5a**) and assessed the mutant clones. Once again, the same escalating doses of rh-TSH were unable to induce proliferation or migration of the cells, regardless of the theoretical increased signal transduction through the TSHR. Moreover, all experiments were conducted under various conditions, such as the presence or absence of insulin (which is theorized to be one of the most important synergistic hormones), the administration of a wide range of doses, the study of different time frames, as well as culture medium replenishments every 24 hours. All these conditions did not produce substantial evidence on the activating role of TSH on the studied cell functions, possibly indicating an inferior role for TSH compared to other parameters.

**Figure 5.**
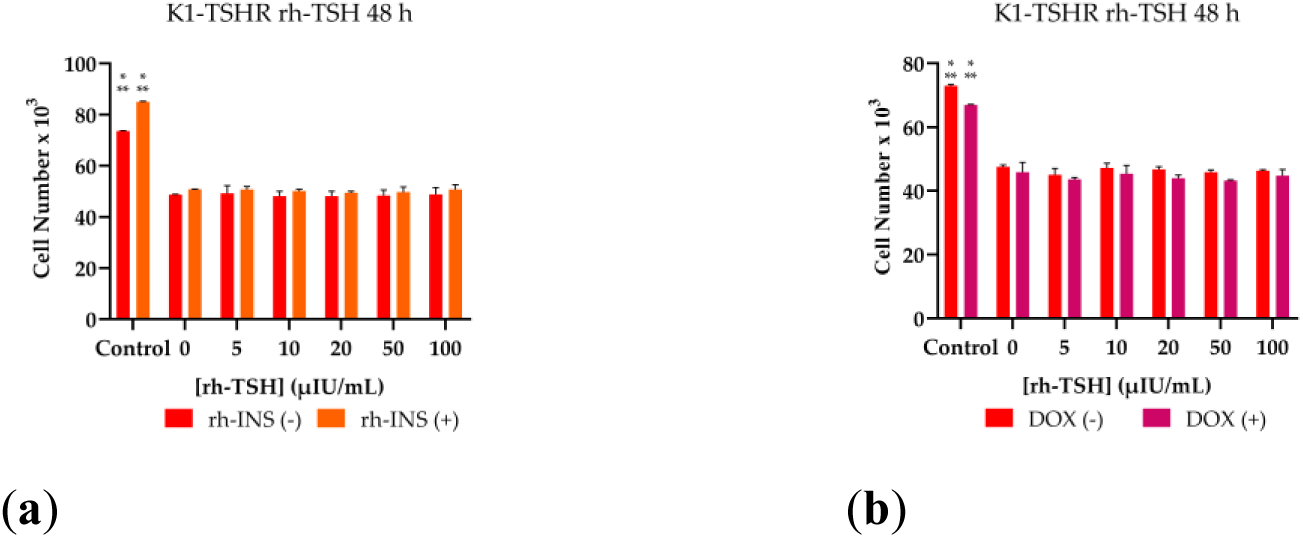
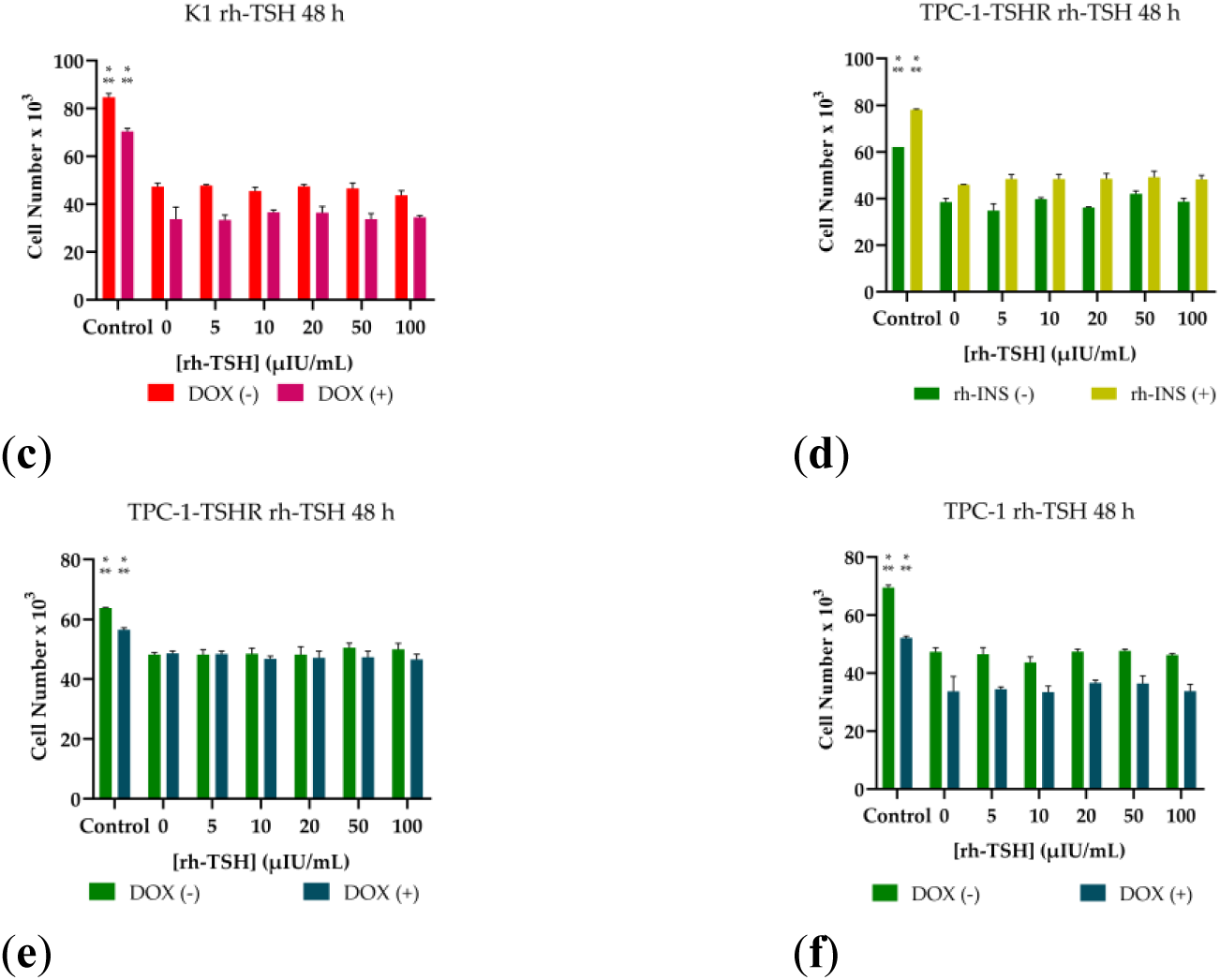
Proliferation Assay of K1-TSHR and TPC-1-TSHR Cells Treated With rh-TSH and/or rh-INS. Transformed cells were seeded into microplates in a medium containing 3 μg/mL doxycycline (DOX) and after 24 hours of settling, attachment and TSHR overexpression, the old medium was aspirated. An assay medium containing 2.5% FBS and the designated doses of the hormones (rh-TSH, h-INS) was added and left for 72 h. A range of rh-TSH concentrations from 5 μIU/mL to 100 μIU/mL was tested in both transformed K1 (K1-TSHR) and TPC-1 (TPC-1-TSHR) cells. **(a)** The red bars represent rh-INS-untreated K1-TSHR cells, while the orange bars represent rh-INS-treated K1-TSHR cells. **(b)** As a control experiment, the effects of rh-TSH were also tested in both DOX-untreated (red bars) and DOX-treated (magenta bars) K1-TSHR cells. **(c)** The effects of DOX were also assessed in (untransformed) K1 cells: the red bars represent DOX-untreated K1 cells, while the purple bars DOX-treated K1 cells. The same set of experiments were performed in TPC-1/TPC-1-TSHR cells **(d)** The dark green bars represent rh-INS-untreated TPC-1-TSHR cells, while the light green bars represent rh-INS-treated TPC-1-TSHR cells. **(e)** DOX-untreated TCP-1-TSHR cells are annotated as green bars while DOX-treated TPC-1-TSHR cells are represented by blue bars. **(f)** The green bars correspond to DOX-untreated (untransformed) TPC-1 cells, while the blue bars correspond to DOX-treated TCP-1 cells. Wherever insulin is annotated (rh-INS), a high concentration of 0.5 U/mL is used. As a positive control for elevated cell proliferation rates, a 10% FBS-containing medium was used which has endogenous b-TSH, b-IGF-II, insulin and other cofactors. All experiments were conducted in triplicate, and the error bars represent the standard error of the mean (SEM). The data analysis was performed using multiple t-tests, and no significant differences between the different rh-TSH concentrations were detected. Insulin-treated cells exhibited an overall increased cell number, which was statistically significant as shown by a paired t-test (p-value<0.001). * Corresponds to a p-value < 0.05; ** corresponds to a p-value < 0.01; *** corresponds to a p-value <0.001.

Although there is a prevailing perception that TSH induces the growth and proliferation of follicular thyroid cells, there is limited evidence as to whether the proliferation is a direct consequence of TSH stimulation, or whether other ligands that act synergistically with TSH activate the TSHR-dependent signaling cascade and lead to observable increased mitotic activity. *In vitro* studies have primarily been conducted using cultures derived from animal cell lines, with only a few studies focusing on human cell cultures that include both normal and malignant thyrocytes. These studies have investigated the effects of various doses of bovine/cattle TSH mainly on cellular responses, particularly regarding cell proliferation. However, the results obtained from these studies have been inconsistent, suggesting that the responses of thyroid cells to TSH treatment can vary widely across different experimental conditions and cell types [19–21,23,26,45–47]. In one study, researchers investigated the effects of escalating doses of b-TSH (1, 10, and 100 mU/mL) on growth and invasion in primary cell cultures of follicular and papillary thyroid carcinomas [45]. Specifically, they analyzed three follicular thyroid carcinoma (FTC) cell lines derived from one patient, comprising one from a primary tumor and two from metachronous metastases originating from a lymph node and lung metastasis. Additionally, they examined two papillary thyroid carcinoma (PTC) cell lines from two patients who had not experienced recurrence. Notably, the maximal effect on proliferation and invasion was observed at a dose of 10 mU/mL (equivalent to 1000x of that found in human plasma), while the dose of 100 mU/mL was found to inhibit proliferation. Intriguingly, the response of the five cell lines varied in terms of growth and invasion at the same b-TSH dose, with PTC lines exhibiting maximal response, followed by FTC lines, and even smaller responses observed in metastatic lines, while inhibition remained consistent [45]. In another *in vitro* study, the authors found that the growth of papillary thyroid cancer cells and adenomatous goiter cells was differentially regulated. Both cell lines were found to express functional TSH receptors. When treated with b-TSH at a dose of 10 μU/mL, a dosage similar to that used in our study, with or without IGF1, the proliferation of cancerous thyrocytes was inhibited, while that of human adenomatous goiter cells was stimulated. Intriguingly, the activation of TK receptors by various growth factors, such as EGF, b-FGF, and IGF1, along with stimulation by E2 (at subclinical concentrations), promoted proliferation in both cancerous and goitrous cells [21]. More recently, a single application of 100, 1000, and 10^4^ μIU/mL of cattle TSH in human PTC cells (TPC-1) increased the proliferation rate in a dose-dependent manner by promoting the transition of TPC-1 cells from the G_1_ phase to the S phase and by remarkably increasing the mRNA and protein expression of cyclin D1 [23]. The contradictory results observed in these various *in vitro* studies could be attributed to several factors, such as the TSH dose used, discrepancies in the specific culture systems used, including variations in cell culture media, supplements, and culture conditions, incubation time, context (normal or cancer cell line), and the method of growth assessment. As FBS contains endogenous TSH (b-TSH), insulin, dissolved iodine salts, and related cofactors that are required for thyroid cell survival and the FBS batches vary significantly, we tried to minimize the effects of these factors by using the minimum required concentration of FBS in our experiments.

It is also noteworthy that most of the published studies have not adequately evaluated the expression of TSHR in the various cell line models. Differences in the media used and multiple passages of cell lines can sometimes have deleterious or silencing effects on the thyroid-like phenotype and the expression of TSHR, which on some phenotypic characterizations have even been reported as absent [42,43]. Many studies even report that important thyroid markers like *TSHR*, *Tg*, thyroid peroxidase (*TPO*), and sodium-iodide-symporter (*NIS*) are not even expressed by the cells, thus creating the question of whether these cells can still be considered a valid thyroid cell model [41]. It is known that the expression of TSHR in primary cultures is 100-fold lower than in intact thyroid cells *in vivo* [48]. To overcome, at least partially, this obstacle, we transformed the two cell lines to overexpress TSHR as a way to secure stable and increased stimulation by TSH. The overexpression was controlled by the presence of doxycycline, and the transformed cells (K1-TSHR and TPC-1-TSHR) were assessed for the same functions as previously noted. Interestingly, no significant changes were observed following stimulation with rh-TSH and insulin, exactly like the non-transformed clones. The migration rates were also unchanged, and the pro-proliferative effects of insulin (once administered alone) were also documented. Our findings are consistent with a prior study wherein 10 mU/mL of b-TSH failed to induce proliferation in human thyroid follicular carcinoma cells that were transfected to overexpress functional TSHR [49].

Besides the expression of TSHR *per se*, it is also important to note the functionality and relevant mutations of the receptor that may affect signal transduction and ultimately cell proliferation. The cell lines that were used in the present work are not known to carry any TSHR mutations, and the TSHR form that is expressed is the wild-type one. The K1 and TPC-1 cell lines are derived from papillary thyroid carcinoma and carry correspondingly the following genetic aberrations: *BRAF p.V600E* and *CCDC6-RET* fusion [50]. Both cell lines have been reported to be genetically identical regarding thyroidal characteristics [51]. Both at baseline and overexpressed levels of wild-type TSHR, we did not see practically any effect on the proliferation of cells descending from rh-TSH stimulation, regardless of the conditions used herein. Additionally, the overexpression of TSHR has been recently tested in TPC-1 cells; however, not in an inducible manner like it is herein presented [40]. In this preprint by Lin et al. in 2022, it is shown that TPC-1 expressed very low levels of the *TSHR* mRNA, and the cell line is transfected with a TSHR-carrying pCDH-CMV-MCS-EF1-Puro (CD510B) plasmid to overexpress the receptor [40]. They report that TSHR overexpression suppressed proliferation; nonetheless, they do not mention any stimulation with TSH (of bovine or human recombinant nature). It is hypothesized that TSHR elevated the phosphorylation levels of IkB, which in turn reduced NF-κB signaling, thus reducing proliferation [40]. However, to this day, no further investigation of the role of overexpressed TSHR has been performed. Moreover, it would be plausible to investigate further the effect of various TSH concentrations not only in wt-TSHR but also in various mutated forms of TSHR that may affect intracellular signaling and thus the proliferation rate, as well as the study of more co-factors.

Further assessment of the signaling pathways downstream of the TSHR, as well as the synergistic actions of other receptors, is also warranted under these conditions and can be the subject of future studies. In order to perform a preliminary investigation of rh-TSH interaction with other circulating hormones, we administered insulin, one of the most important hormones that acts synergistically with TSH [26,52–54]. Besides insulin, insulin-like growth factors (IGFs) are parts of a protein family with many roles in both physiological functions and disease [55,56]. Both IGFs and insulin are known to bind the insulin receptor (IR) as well as the insulin-like growth factor receptors I and II (IGF1R, IGF2R), which can dimerize with the TSHR and transduce signals. Insulin has a lower affinity for the IGF1R compared to IGF1; however, higher concentrations of insulin have been found to successfully activate the receptor [57–59]. Specifically, the insulin-like growth factor I receptor (IGF1R) is a transmembrane tyrosine kinase receptor involved in growth and development [60]. It seems that there is a cross-talk between the TSH/TSHR and insulin/IGF1/IGF1R pathways in thyroid cell proliferation and differentiation [53,54,61–66]. FBS contains both bovine IGF1 and bovine insulin, both of which are adequate to maintain the normal growth of the cell lines since they are equipotent to the human hormones [67,68]; however, in an attempt to overstimulate the IGF1R which has a significant crosstalk with TSHR, we modified the culture medium to contain a high concentration of recombinant human insulin (Humalog®). In agreement with our experimental design and data, in most of the *in vitro* studies, IGF1 or high insulin levels are required for the culture to achieve the receptor’s activation [26,69]. According to our experiments, the addition of insulin at a concentration of 0.5 U/mL in the medium on top of rh-TSH increased the cell number, but not in a rh-TSH dose-dependent manner.

TSH, besides proliferation, modulates thyroid hormone synthesis and secretion by regulating the expression of various thyroid-specific genes such as *TSHR*, *Tg*, *TPO*, and *NIS*, all of which are considered markers of thyroid cell phenotype, and their maintenance guarantees that no de-differentiation has occurred. Our findings indicated that treatment of K1 and TPC-1 papillary thyroid cancer cells with increasing concentrations of rh-TSH led to a gradual increase in expression of the *Tg* gene, with peak mRNA expression levels at the highest dose. Our results seem to be consistent with previous studies where treatment of cultured thyroid cells with b-TSH stimulated the *TPO* and *Tg* mRNA levels in a dose- and time-dependent manner [70,71]. A limitation of our study is that it is based on *in vitro* experiments and employs cell lines that may gradually lose some of the properties of the original thyroid tissue as a result of mutations and/or epigenetic silencing resulting from selection pressure during prolonged cell culture [42,72]. To at least partially address this concern, we evaluated our cell lines for the preservation of thyroidal phenotype by assessing *Tg* mRNA expression, observing that the cells retained some features of their original phenotype since the signaling pathways (and enzymes) that regulate Tg synthesis are specific to thyroid cells. However, it is noteworthy that under basal conditions, *TSHR* mRNA expression was low in our cancerous cell lines, while normal thyroid tissue has significantly higher expression of the receptor. To overcome this limitation, we engineered both cell lines to overexpress the thyrotropin receptor. To strengthen our findings, further investigations are warranted using additional cell types that overexpress wild-type as well as mutant *TSHR* variants (various versions of constitutively active or inactive, as aforementioned) [36] or utilizing more physiologically relevant cell systems such as primary cultures of thyroid gland organoids [73] from cancerous thyroid tissue. Additionally, it is imperative in future studies to assess the functionality of the overexpressed TSHR by assessing the activation status of downstream signaling pathways that were not assessed in this study, given that no effect was seen in proliferation, and possibly expand our understanding of the synergistic roles of IGFs and their receptors.

## 5 Conclusions

In conclusion, we have shown that rh-TSH in clinically relevant doses cannot induce proliferation and migration in papillary cancerous cell lines under various conditions. Further research is warranted to dissect the molecular mechanisms underlying these effects. Our results could translate into better management of PTC patients in the future.

## 7 Author Contributions

Conceptualization, M.M. and P.K.; methodology, G.K., A.V., C.A., I.M., and A.S.; software, G.K. and A.S.; validation, G.K., A.V., and C.A.; formal analysis, G.K., C.A., and A.S.; investigation, G.K., A.V., C.A., and I.M.; resources, M.M., D.C., and P.K.; data curation, G.K.; writing—original draft preparation, G.K., D.C., and M.M; writing—review and editing, G.K., D.C., M.M, and P.K..; visualization, G.K.; supervision, P.K.; project administration, M.M and P.K..; funding acquisition, M.M.

All authors have read and agreed to the published version of the manuscript.

## Acknowledgments

We kindly thank Dr. Holger Jaeschke from University Hospital Essen for providing us with the plasmid used during our cloning experiments. We also thank Dr. Konstantina Nika from the School of Medicine of the University of Patras for her aid in the cloning experiments and providing us with resources.

## 9 Data Availability Statement

We confirm that the data supporting the findings of this research are contained within the article and its supporting material.

## 10 Appendices

### Appendix A

In this part, the procedure that led to the use of 2.5% FBS in the proliferation ans migration assays is described. FBS is vital for cell culture; however, it contains b-TSH, b-INS, and b-IGFs that would interfere with the assays. Since its omission was unfeasible, we cultured the cells with different FBS concentrations to settle on the least concentration that would adequately support proliferation for 72 h without having excessive amounts of the hormones. The cells were found to thrive at the 10% and 5% FBS concentrations; nevertheless, dropping the FBS percentage to less than 2.5% was leading to significantly reduced proliferation following 72 h of culture (**Figure A1**). Therefore, we regulated the FBS concentration at 2.5% for all assays, while the baseline culture conditions remained stable at 10% FBS to avoid thyroid cell de-differentiation due to a lack of proper stimulation from b-TSH, b-INS, b-IGF1, and sufficient iodine.

**Figure A1.**
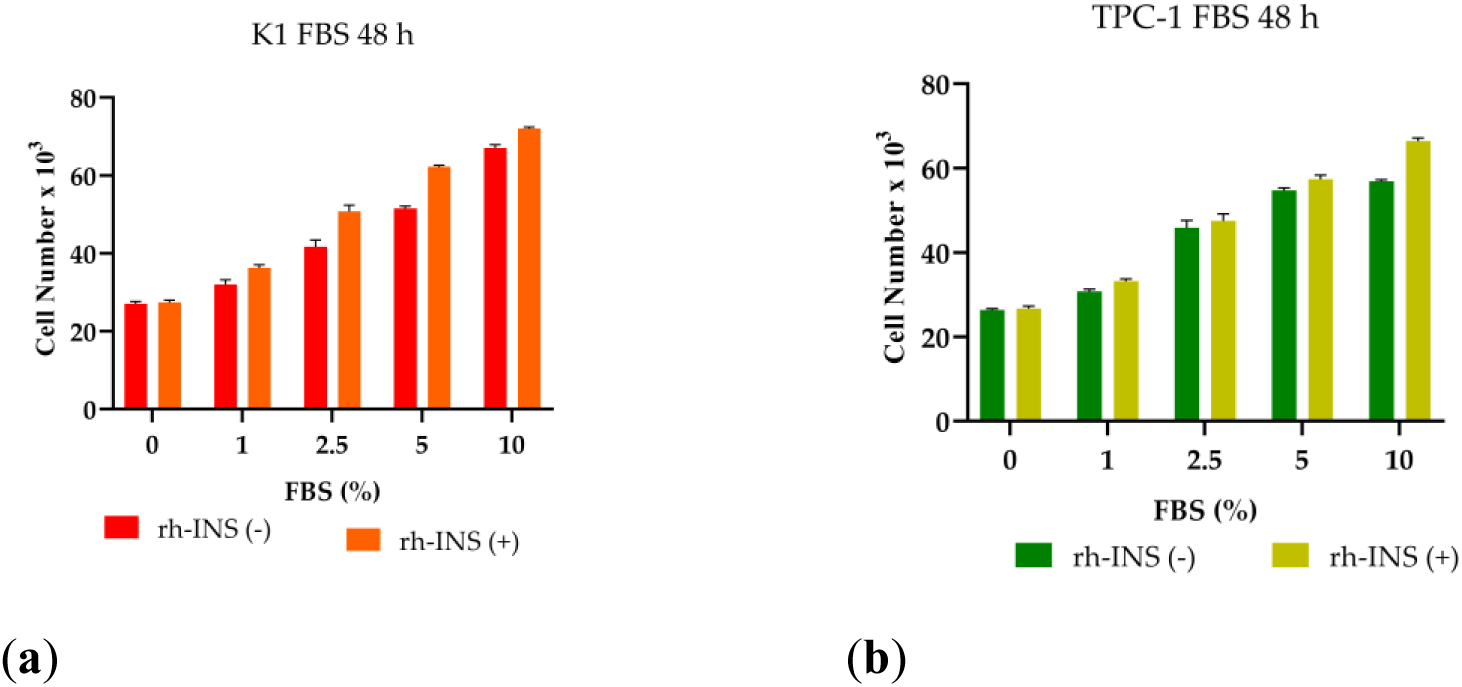
Minimal FBS Concentration Optimization. (**a**) K1 cells were seeded into microplates, and after 24 hours of settling and attachment, the old medium was aspirated and replaced with a new one containing different doses of FBS (2.5%, 5%, and 10%). The 2.5% FBS concentration allowed the cells to survive for 72 h, while the final cell number was significantly lower than that observed when greater doses were used. Therefore, it was designated as the standard medium for all the experiments. The red bars represent rh-INS-untreated K1 cells while the orange bars represent rh-INS-treated K1 cells. (**b**) The same effect was observed on the TPC-1 cells. The dark green bars correspond to rh-INS-untreated TPC-1 cells while the light green bars represent rh-INS-treated cells. The experiments were conducted in triplicate, and the error bars represent the standard error of the mean (SEM).

### Appendix B

Herein, supportive experiments that were performed to validate the successful transformation of the cells are presented. First, the creation of plasmids containing the *TSHR* gene was verified by isolating plasmids from transfected *E. coli* and analyzing them using agarose gel electrophoresis. As shown in the gel image from the transilluminator, the predicted size of 2.5 kb was detected, thus verifying the successful construction (**Figure A2a**). Afterwards, the isolated plasmids were used to chemically transfect HEK293 cells to produce lentiviruses containing the Lenti-X Tet-on-Advanced System and the *wt-TSHR* gene. The lentiviruses were used to transduce K1 and TPC-1 cells, which were subsequently selected using puromycin and geneticin (G418 sulfate). The transfected cells were assessed for successful transformation by isolating the total DNA and detecting the insert using PCR (**Figure A2b**). The primers for the newly inserted *TSHR* cDNA were designed to avoid detecting the genomic DNA (gDNA) *TSHR* alleles. Both cell lines were found to have integrated the inserts; therefore, the transformed cells were named K1-TSHR and TPC-1-TSHR, respectively.

**Figure A2.**
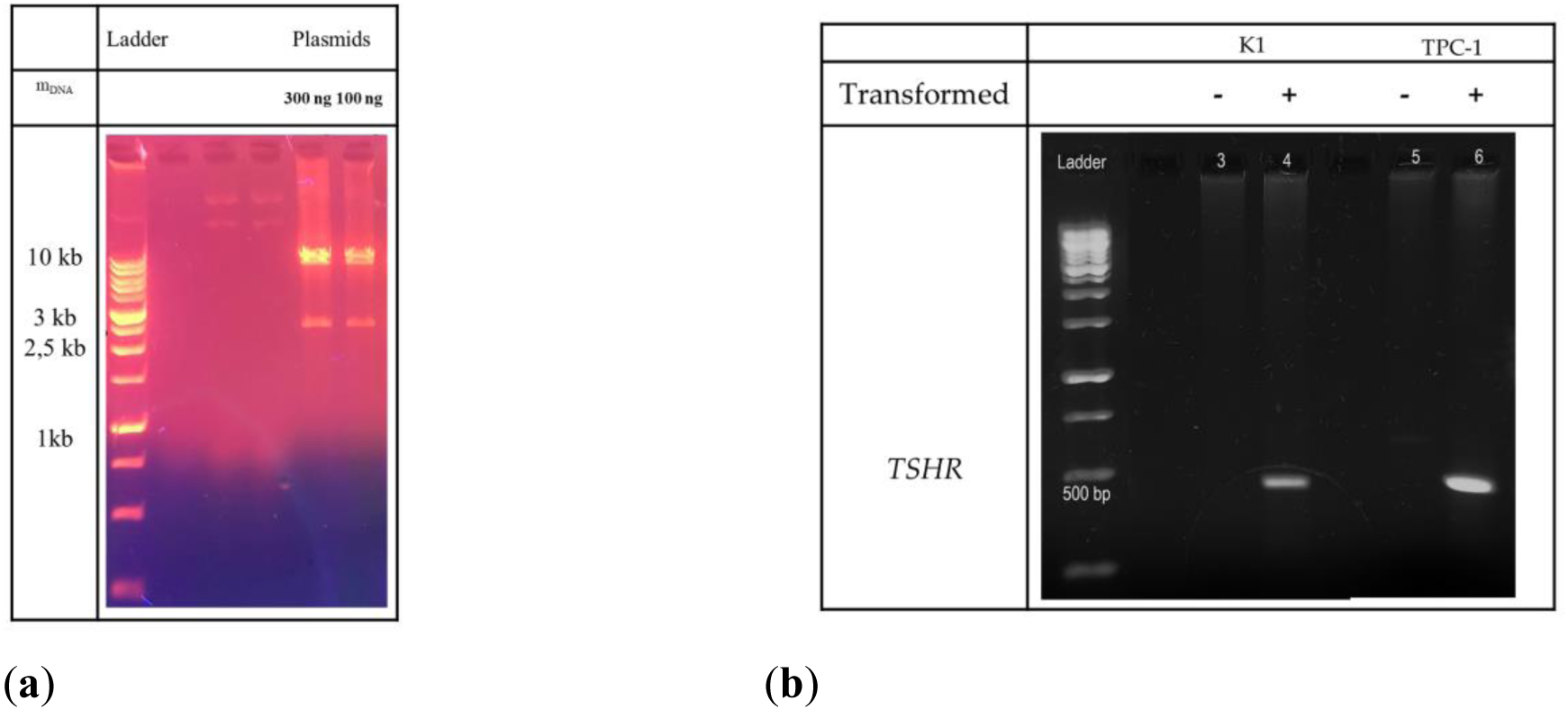
Successful Plasmid Construction and Cell Transfection Verification. (**a**) *wtTSHR* cDNA was inserted into pLVX-Tight-Puro plasmid in EcoRI and BamHI positions. The expected plasmid had a predicted size of 2.5 kbp and was successfully detected. Different quantities are displayed (300 μg and 100 μg of DNA at lane five and lane six, respectively); (**b**) After the transduction of the K1 cells (lane two), and the TPC-1 cells (lane four) with lentiviruses and the selection using the antibiotics, the survived cells were lysed, and total DNA was isolated using the Monarch® Genomic DNA Purification Kit by NEB. PCR was conducted and revealed that the transformed clones had successfully incorporated the *TSHR* cDNA. The expected PCR product had an expected size of 439 bp. As a control sample, the non-transformed cells were used (lanes 1 and 3, respectively).

### Appendix C

In this part, a full matrix of all screened rh-TSH doses (plus combinations with insulin) is presented.

**Figure A3.**
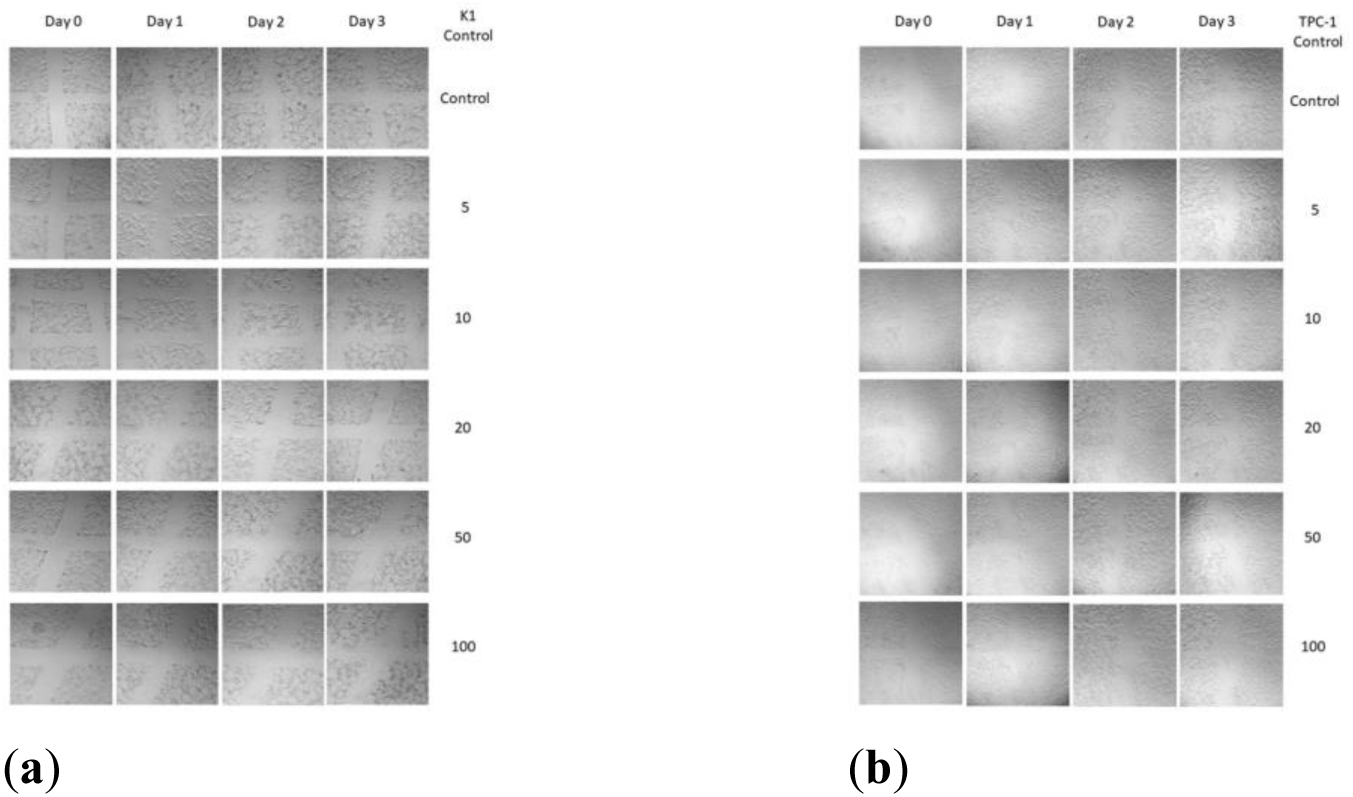
Wound Healing Assay of K1 and TPC-1 Cells. Cells were seeded into six-well microplates, and after the formation of a confluent monolayer, scratches of equal dimensions were made using a pipette tip. Subsequently, the old medium and the debris were aspirated and replaced with a new one containing 2.5% FBS, a range of rh-TSH doses from 5 μIU/mL to 100 μIU/mL and 0.5 U/mL rh-INS. Photographs were taken every 24 hours using a camera mounted on a microscope (100X magnification). No changes were observed in either non-transformed cells (**a**) or transformed cells (**b**).

**Figure A4.**
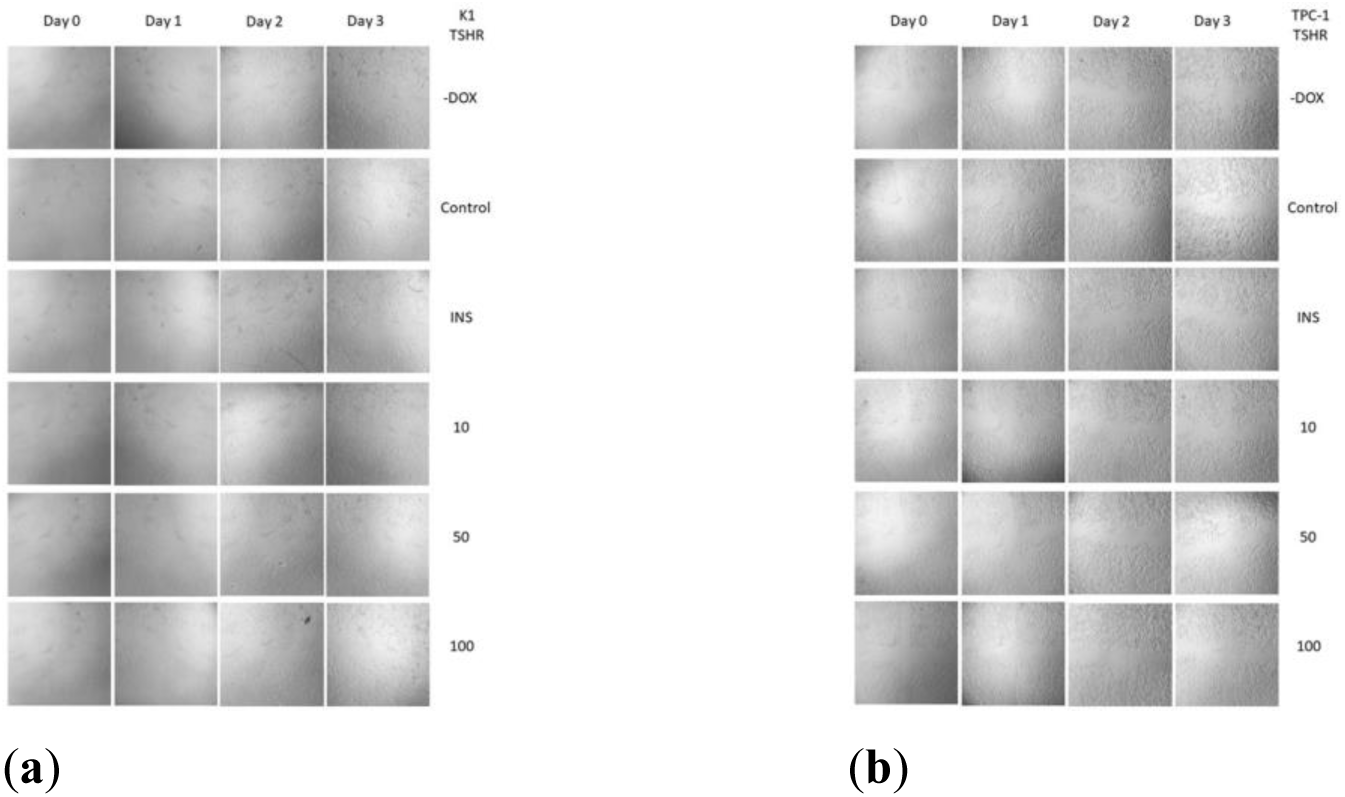
Wound Healing Assay of K1-TSHR and TPC-1-TSHR Cells. Cells were seeded into six-well microplates, and after the formation of a confluent monolayer, scratches of equal dimensions were made using a pipette tip. Subsequently, the old medium and the debris were aspirated and replaced with a new one containing 2.5% FBS, a range of rh-TSH doses from 5 μIU/mL to 100 μIU/mL and 0.5 U/mL rh-INS. Photographs were taken every 24 hours using a camera mounted on a microscope (100X magnification). No changes were observed in either non-transformed cells (**a**) or transformed cells (**b**).

